# Plant defense metabolite detoxification by a pathogenic fungus opens necrotic tissue to diverse bacteria in *Arabidopsis thaliana*

**DOI:** 10.1101/2025.08.22.671692

**Authors:** Shubhangi Sharma, Jingyuan Chen, Matthew Agler

## Abstract

Foliar necrotic fungal pathogens pose a constant threat to plant health. Their virulence can be affected by both the host immune system, as well as diverse host-associated bacteria. Generally, how these factors interact is not well studied, but may provide insights toward sustainable plant protection. *Sclerotinia sclerotiorum* (Ssc) is a devastating, broad host range necrotrophic pathogen with limited control options. Here, we find that necrotic Ssc lesions strongly enrich diverse bacterial taxa. *Arabidopsis thaliana*, like other Brassica plants including crops, produce and store large amounts of aliphatic glucosinolates (aGLS). Upon Ssc invasion and tissue damage, myrosinase enzymes transform aGLS into toxic metabolites, including aliphatic isothiocyanates (aITCs). Adapted Ssc strains can detoxify aITCs by hydrolyzing them with the enzyme SaxA, which is important for its virulence. Here, we find that this detoxification is critical for bacterial enrichment in necrotic lesions, since a SaxA-deficient strain does not alter the bacteriome. Additionally, our results suggest that both aliphatic and other GLSs likely contribute specific effects to this phenomenon. Thus, we demonstrate a mechanism in which the interaction of host and pathogen factors together shape a pathogen-associated bacteriome, providing directions to better understand interactions of host bacteria with foliar necrotic pathogens.

## 1. Introduction

Most plant tissues are colonized by a variety of microorganisms, including bacteria, fungi, and other eukaryotes that play critical roles in plant health and productivity. Bacterial diversity is of particular interest because of the direct and indirect ways it protects plants, for example from detrimental environmental pathogens (Duran et al., 2018; Paasch et al., 2023). In leaves, bacterial diversity is largely controlled by filtration effects exerted by plant host factors and the environment. (Wagner et al., 2016). Importantly, these factors can act strongly via inter-kingdom microbe-microbe interactions (Agler et al., 2016). For example, foliar fungal biotrophic pathogens cause strong changes to leaf bacterial community diversity upon infection by increasing the abundance of diverse bacterial taxa (Duran et al., 2021; Jakuschkin et al., 2016). Bacteria recruited to leaves during pathogen infection can influence disease outcome (Goossens et al., 2023), so it is important to understand these processes. It has been suggested that rearrangements of host defense and metabolism by pathogens could play an important role, but the mechanisms have in most cases not yet been uncovered.

Plant specialized metabolites play important roles in plant protection especially against herbivores and nonhost pathogens. A well-studied defense system in the Brassicaceae, including important crops and the model plant *Arabidopsis thaliana* is the aliphatic glucosinolate (aGLS) - myrosinase system. aGLSs are amino acid-derived secondary defense metabolites which are not by themselves biologically active and are stored intact at high levels within the vacuoles of specialised leaf cells. Upon tissue damage, they come into contact with myrosinase enzymes (β-thioglucoside glucohydrolase), and are rapidly broken down into diverse toxic metabolites, commonly known as the “mustard oil bomb” (Halkier and Gershenzon, 2006). In the model *A. thaliana* genotype Col-0, the main breakdown products are aliphatic isothiocyanates (aITCs). These highly reactive compounds are well-recognised for their antimicrobial activity against both bacterial and fungal plant pathogens (Chen et al., 2020; Fan et al., 2011).

Host-adapted bacterial and fungal pathogens of *A. thaliana* express ITC hydrolases (metallo-β-lactamase enzymes) known as SaxA (“<u>s</u>urvival in <u>A</u>rabidopsis e<u>x</u>tracts”) enzymes. These help them handle the massive release of ITCs upon tissue destruction by hydrolyzing them to non-toxic amines. Pathogens without these hydrolases are compromised in virulence in *A. thaliana* (Chen et al., 2020; Fan et al., 2011). Besides pathogens, ITCs like sulforaphane in the *A. thaliana* genotype Col-0 are also toxic to bacteria that colonize healthy plant leaves. Indeed, a subset of these bacteria express similar activity that can detoxify ITCs (Unger et al., 2024). Thus, both bacterial and fungal pathogens modify the *A. thaliana* leaf chemical environment upon invasion by instigating glucosinolate hydrolysis into toxic products, as well as by detoxifying those products.

*Sclerotinia sclerotiorum* de Bary (Ssc) is a highly destructive fungal necrotrophic pathogen with a wide host range, including important Brassicaceae crop plants. It expresses a SaxA protein that increases its virulence on *A. thaliana* by detoxifying GLS breakdown products. Little is known about how Ssc influences the leaf microbiome or the role of ITCs in this process. We hypothesized that during leaf necrosis, ITC release and its subsequent hydrolysis would modulate assembly of a pathogen-associated bacterial community. To test this hypothesis, we studied the effect of Ssc infection on local and systemic leaf bacteriomes of *A. thaliana* Col-0. To understand how aGLSs and their breakdown products underlie observed effects, we looked at bacteriomes in the Col-0 WT and its aGLS-free *myb28/myb29* mutant (Sønderby et al., 2007) infected with the necrotrophic fungus *S. sclerotiorum* (Ssc WT) or the Ssc Δ*SaxA* mutant (Chen et al., 2020), which cannot hydrolyse ITCs.

## 2. Materials and Methods

### 2.1 Fungal strains and growth conditions

The *Sclerotinia sclerotiorum* strain 1980 wild-type (Ssc WT) and the knockout mutant strain Ssc Δ*SaxA* (Chen et al., 2020) were routinely cultured on potato dextrose agar (PDA) at 24°C for 4 days. In order to prepare fungal inoculum, the fungus was cultured in liquid potato dextrose broth (PDB) at 24°C and gentle shaking at 120 rpm for 3 days. The fungal biomass was strained in a sterile nylon mesh and washed with autoclaved miliQ water until the flow-through was clear to remove all PDB. The washed fungal biomass was crushed in a blender along with fresh 40 mL of PDB at the slowest speed for 40s. The number of fungal hyphal fragments were counted in a Thoma cell counting chamber and the fungal inoculum was adjusted to 2×10^5^ fragments per mL using PDB.

### 2.2 Plant cultivation, inoculation and sampling

*Arabidopsis thaliana* ecotype Columbia (Col-0) and the *myb28/29* double knockout mutant (in the Col-0 background) which does not produce aliphatic GLS was used. Seeds were vernalised in 0.1% agarose at 4°C for 5 days. A standard laboratory potting soil was made by mixing 4L of Floraton 3 potting soil (Floragard), 2L perlite (0-6mm Perlite Perligran, Knob), and 25g Substral Osmocote garden flower fertilizer (Celaflor). This soil was supplemented with a soil slurry inoculum made by dissolving 15g of dried garden soil (collected from a common garden outside our laboratory) in 100 ml of miliQ water. Two parts soil slurry was used to wet four parts of dry laboratory potting soil. The plants were grown on an LED light shelf under 14-h days at about 260 μmol·m^−2^ ·s^−1^ for four weeks.

### 2.3 Ssc inoculation and sampling

Four-week old *A. thaliana* Col-0 or *myb28/29* plants were infected with *S. sclerotinia* (WT or Δ*SsSaxA*) hyphal suspension or inoculated with a PDB only control (Figure 1A). Ten individual plants of each genotype were randomly assigned to each treatment. Five leaves per plant were inoculated with a 5µl drop. The plants were covered with a lid to increase humidity for successful infection. After development of symptoms (36-38h, (Figure 1B)), three tissues were taken from each plant: The necrotic zone of the leaf constituting the infected leaf zone (ILZ), the remaining leaf tissue constituting the non-infected leaf zone (NILZ), and non-inoculated leaves (NIL) as systemic tissue. Each sample consists of combined tissues collected from three leaves from a single plant. The PDB control plants were treated the same, but the “ILZ” corresponds to the spot where PDB was inoculated and there was no necrotic tissue **(Figure 1C)**. All samples were flash-frozen using liquid nitrogen and stored at −80°C.

**Figure 1:**
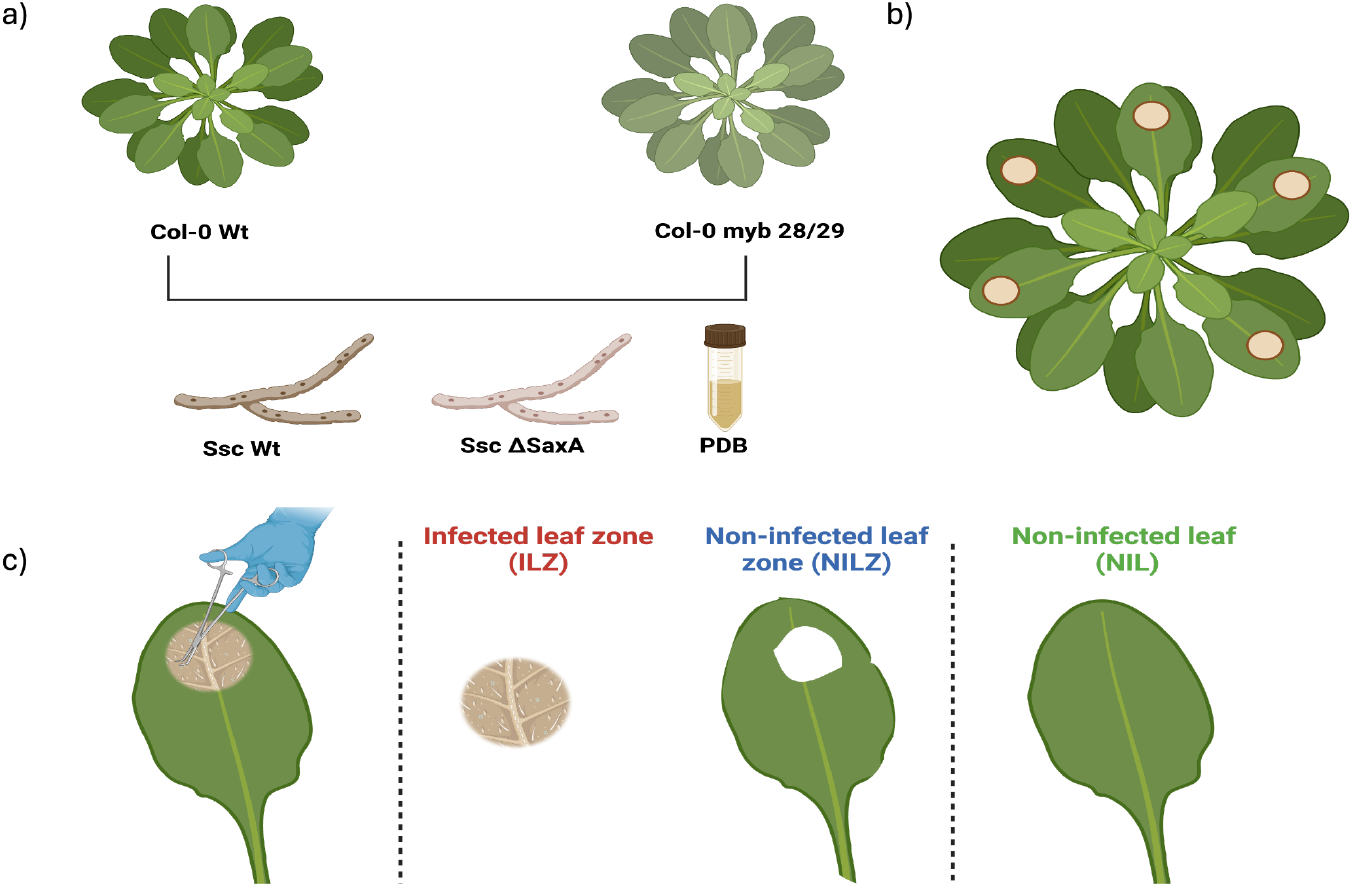
Inoculation and sampling strategy for *S. sclerotiorum* infected *A. thaliana* leaves. A) A. thaliana Col-0 and myb28/29 mutant plants grown in a natural soil-supplemented substrate were drop-inoculated with Ssc WT, Ssc Δ*SaxA* or a PDB control. B) Leaves inoculated with Ssc developed necrotic lesions at the site of inoculation. C) Inoculated leaves were dissected so that bacterial communities could be characterized in necrotic tissue (ILZ), non-necrotic tissue of the same leaf (NILZ) and in systemic tissue (NIL). PDB-inoculated control plants were dissected similarly. A more detailed version of the figure is provided in Figure S1.

### 2.4 DNA extraction and 16S rRNA gene amplicon library preparation and sequencing

DNA was extracted from all the samples using a well-established method. Briefly, the frozen plant tissue was lysed using a bead beater for 30 s at 1,400 rpm. To the lysed plant tissue, 150 µL of 2x cetyl trimethylammonium bromide (CTAB) buffer (2% cetyltrimethylammonium bromide, 1% polyvinylpyrrolidone, 100 mM Tris-HCl, 1.4 M NaCl, 20 mM EDTA) was added followed by a ten-minute incubation at 65°C. The supernatant from the CTAB extraction was mixed with 200μL of ice-cold phenol-chloroform-isoamylalcohol (25:24:1), centrifuged at top speed for 5 minutes, then the supernatant was precipitated overnight at −20C in an equal volume of isopropanol. After removing the isopropanol, the pellet was washed twice with 80% ethanol, air dried and resuspended in Tris-HCl (10mM, pH 8).

The extracted DNA was used as a template in a 2-step 16S rRNA gene amplicon sequencing protocol modified to quantify bacterial loads via host-associated microbe PCR (hamPCR) (Lundberg et al., 2021). In short, 16S amplification was performed together with amplification of a single copy host plant gene (GI = *GIGANTEA* gene, referred to as “GI gene”). ZymoBIOMICS Microbial Community DNA Standard II (ZymoResearch, Freiburg, Germany; referred to as “ZymoMix”) was used as positive control in three reactions, and eight negative controls included a no template PCR control, two reactions with CTAB DNA extraction buffer, two reactions with TRIS buffer that was used to resuspend DNA, and three reactions with nuclease free water. In a first 5-cycle PCR, samples were amplified using 341 F/799 R “universal” 16S rRNA primers and GI primers, both modified with an overhang sequence. The reaction included blocking oligos to reduce plastid 16S amplification (Mayer et al., 2021). The PCR product was enzymatically cleaned and used as template in a second, 35-cycle, PCR to add index barcodes and sequencing adapters using primers that bound to the overhang region. PCR products were cleaned up with magnetic beads and libraries were quantified using PicoGreen (1:200 diluted stock, Quant-iT™ PicoGreen™, ThermoFisher) in a qPCR machine (qTower3, JenaAnalytik, Jena, Germany). Samples were pooled according to their normalized fluorescence relative to the highest fluorescent sample. The pool was further processed to increase the fraction of 16S relative to GI, as recommended in the original protocol. Libraries were sequenced on an Illumina MiSeq instrument for 600 cycles. PCR reaction details and primer sequences are previously described in (Unger et al., 2024).

### 2.4 Data analysis and statistical methods

The amplicon sequencing data was split into samples based on indices and adapter sequence trimmed from distal read ends using Cutadapt 3.5 (Martin, 2011). We then clustered amplicon sequencing data (forward reads only as they were much higher quality) into amplicon sequencing variants (ASVs) using dada2 (Callahan et al., 2016). Chimeric sequences were removed and a sequence table retrieved from the data. Taxonomy was assigned to the final set of ASVs using the Silva 16S rRNA (v 138.1) database (Quast et al., 2013). The database was supplemented by adding the *A. thaliana* GI gene sequence.

Further filtering and quality control was performed using the R package phyloseq (McMurdie and Holmes, 2013). First, the data was filtered to remove reads derived from mitochondrial or chloroplast 16S, then was split into bacterial 16S and host GI gene sets. The positive and negative controls were then checked. The positive controls looked as expected. After removing ASVs derived from the positive controls, of 11 controls (three positive controls and eight negative: one no template PCR control, two with the TRIS buffer that was used to resuspend DNA, two with CTAB DNA extraction buffer that was carried through the process and three nuclease free water controls), one positive control and one TRIS negative control contained contamination. Thus, only a small amount of contamination is expected in the samples and given the replication the results should still be very robust. Based on this analysis, we decided to keep samples with at least 100 bacterial reads (the average total read depth was ~14000 reads/sample and most had much more than 100 bacterial reads, but as expected for leaf samples, samples with low bacterial loads contain many plant chloroplast and mitochondria reads that get filtered out).

To generate each figure, the quality controlled data was first subset to the samples of interest. All analyses were performed after grouping the ASVs at the genus level. Alpha diversity parameters were measured using the function “estimate_richness” and plotted in a boxplot. A t-test was used to test for significant differences. For beta diversity analyses, the abundance data was first adjusted for the number of host GI reads to reflect absolute abundances. Next, bray-curtis distances between samples were calculated with the function “distance” and a constrained analysis of principle coordinates was performed with function “ordinate”, constrained for the tissue type (ILZ, NILZ or NIL). A PERMANOVA analysis was performed using the function “adonis2” to evaluate significance of the tissue type on the bray-curtis distances with 1000 permutations. Finally, the ordination was plotted using the function “plot_ordination”. An estimate of bacterial load was calculated for each sample by dividing the sum of bacterial reads by the bacterial reads + GI reads. These were square-root transformed before plotting and a t-test was used to check for significance between treatments. To test for differential abundance of the bacterial taxa between leaf tissues for a given treatment, we used DESeq2 (Love et al., 2014). Because some treatments had far more significant taxa than others, the cutoff p-value was lowered for these treatments to only show a reasonable number of taxa. The p-values used are described in the corresponding supplementary figures. The log-transformed abundances of taxa identified by DESeq at the p-value cutoff were plotted in boxplots for visual inspection. For the plots we additionally included a p-value based on a non-parametric Kruskal-Wallis test, as we occasionally observe that taxa identified by DESeq have insignificant differences between treatments.

## 3. Results

### 3.1 *S. sclerotinia* infection alters the leaf bacteriome by enriching diverse bacteria in necrotic tissue

To test whether disease caused by the necrotrophic fungal pathogen *S. sclerotinia* shapes bacterial communities inhabiting *A. thaliana* Col-0 leaves, we infected five leaves of four-week-old plants with hyphal fragments of Ssc WT suspended in PDB medium or with only PDB as a control (Figure 1). Infection of Col-0 WT by Ssc WT increased the estimated total bacterial diversity in necrotic tissue (the ILZ) compared to PDB treatment (ILZ, Chao1 p = 0.012) but did not affect the evenness in the community (Shannon Index, p = 0.5) (Figure 2A and 2B). Additionally, the ILZ of infected plants exhibited a significantly increased bacterial load (16S/GI p = 0.033) (Figure 2C). In comparison, there was no effect on diversity or bacterial loads in neither the non-infected part of the treated leaf (NILZ) nor in a systemic healthy leaf (NIL). Similarly, the Ssc WT treatment resulted in a significant shift in the bacterial community structure in the ILZ, with tissue type explaining about 24% of variation in the bacterial community (Figure 2E, PERMAVONA, p = 0.001). The control PDB treatment had no significant effect (Figure 2D, PERMAVONA, p = 0.123). DESeq2 differential abundance analysis identified numerous bacterial genera whose abundance differed between the ILZ and the NILZ. Specifically, 25 genera across Proteobacteria, Actinobacteriota and Bacteroidota were more abundant in the ILZ, while none were more abundant in the NILZ (with fdr-adjusted p < 0.001). Most of the taxa were highly enriched: All had a log2FC of at least 4.5 and all but one were differentially enriched with p<0.05 based on a subsequent Kruskal-Wallis test (Figure S2 and Supplementary Table 1). In contrast, PDB treatment resulted in 12 differentially enriched genera, all higher abundance in the NILZ (p < 0.05 - the cutoff is higher because otherwise no taxa are found). The suppressive effect of PDB treatment was not strong - only two of the 12 genera were differentially enriched based on a subsequent Kruskal-Wallis test (Figure S3). Collectively, these results are consistent with leaf infection by the fungal pathogen Ssc resulting in strong and even enrichment of diverse bacterial taxa in the necrotic ILZ.

**Figure 2:**
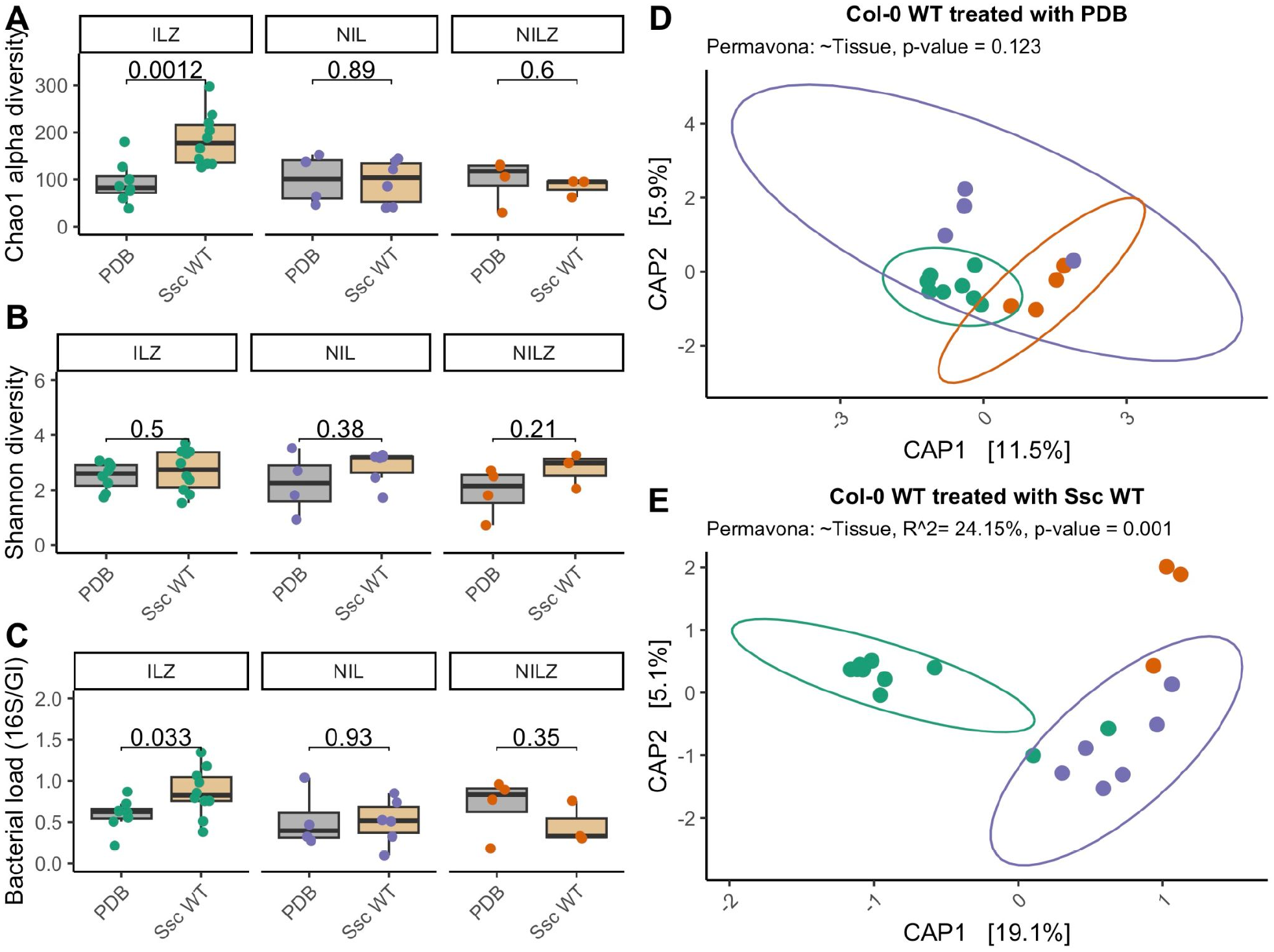
Bacterial community analysis of *A. thaliana* Col-0 WT infected with *S. sclerotiorum* WT. Measurements of leaf bacterial community composition of different leaf tissues (ILZ, NILZ and NIL - Defined in Figure 1) infected with Ssc WT. Alpha diversity results are based on 16S rRNA gene amplicon sequencing data, grouped at the genus level, while load and beta diversity results are the same data scaled to reflect absolute abundance (hamPCR, see methods). Each point represents one leaf sample and is colored according to the leaf tissue: Green: ILZ, Purple: NIL, Orange, NILZ. A) Chao1 estimate of total alpha diversity, B) Shannon Index and C) Bacterial load in tissues from Ssc-infected plants (tan bars) compared to the control PDB treatment (grey bars). P-values are the result of a t-test comparing the treatments. D) and E) Constrained analysis of principle coordinates showing the effect of tissue type on the bacterial community structure, based on bray-curtis distances. PERMANOVA values reflect the result of 1000 permutations of the data.

### 3.2 Fungal ITC degradation shapes bacterial enrichment in the Ssc necrotic zone

To assess how fungal ITC degradation (SaxA activity) contributes to altering the bacterial community in leaves infected with the fungal pathogen Ssc, we next infected *A. thaliana* Col-0 genotype plants with the Ssc Δ*SaxA* mutant. This strain is impaired in the detoxification of ITCs. The experimental setup was identical to before. After infection with Ssc Δ*SaxA*, only a tendency for an increase in estimated total bacterial diversity was observed in the ILZ (Chao1 p = 0.068) with no effect on systemic tissues (NILZ or NIL) (Figure 3A and 3B). The bacterial load, on the other hand, was still strongly increased in the ILZ of Ssc Δ*SaxA*-infected leaves (Figure 3C, p = 0.0004). Despite these changes, Ssc Δ*SaxA* infection did not result in a more significant shift in beta diversity than PDB treatment (Figure 3D and 3E, PERMANOVA, p=0.125). The overall weaker effects can be explained by a much more limited number of genera enriched in the ILZ (with fdr-adjusted p < 0.001). Only five genera were enriched (Figure S4 and Supplementary Table 1), all of which were also enriched in the ILZ during Ssc WT infection (Figure S3). Together, necrotic tissue infected by the Ssc Δ*SaxA* mutant enriched a small subset of the bacteria that Ssc WT lesions enriched, suggesting that ITC detoxification is required to open the necrotic lesions to a wide bacterial diversity.

**Figure 3:**
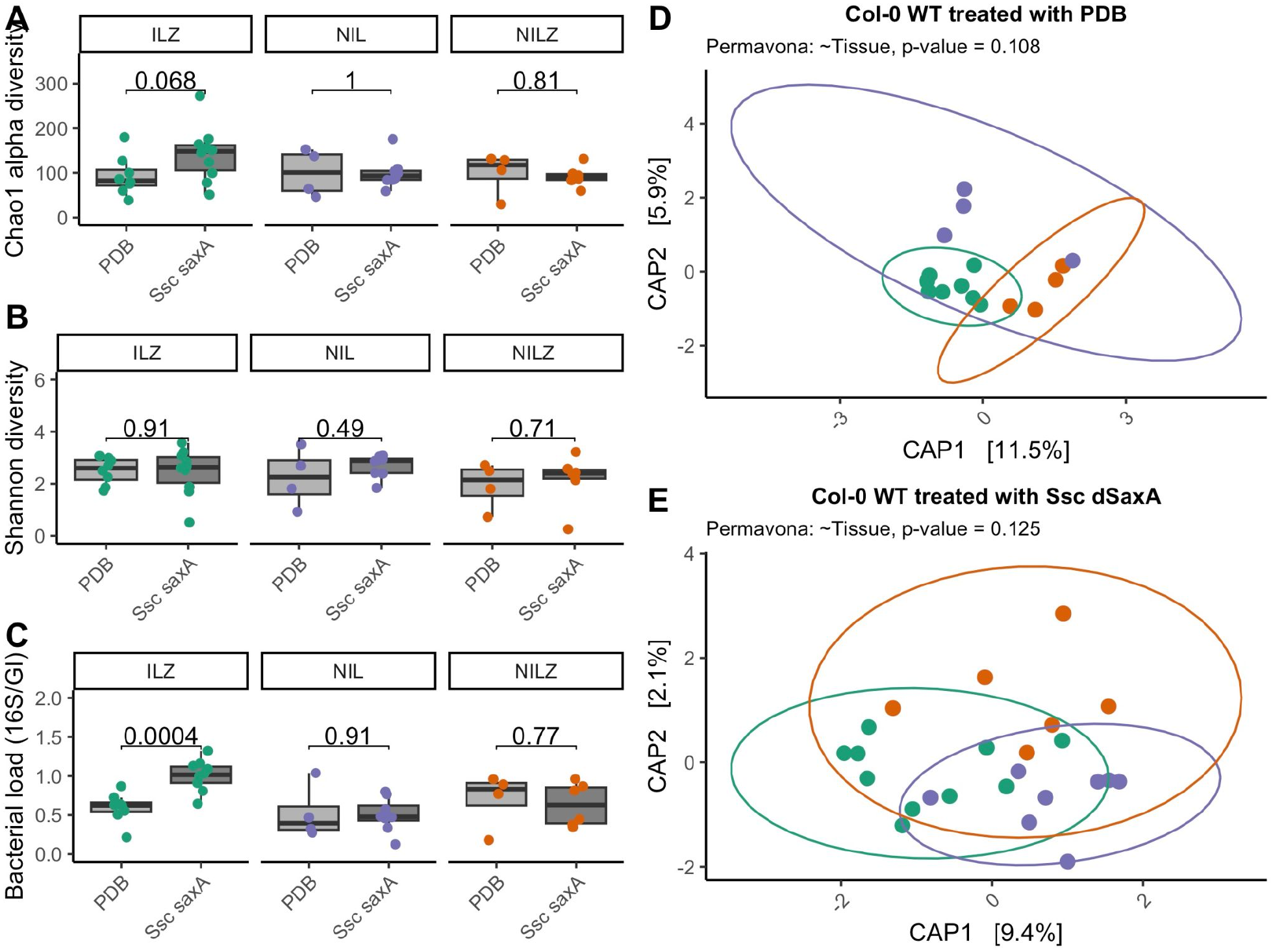
Bacterial community analysis of *A. thaliana* Col-0 WT infected with the *S. sclerotiorum* Δ*SaxA* mutant. Measurements of leaf bacterial community composition of different leaf tissues (ILZ, NILZ and NIL - Defined in Figure 1) infected with Ssc Δ*SaxA* mutant. Alpha diversity results are based on 16S rRNA gene amplicon sequencing data, grouped at the genus level, while load and beta diversity results are the same data scaled to reflect absolute abundance (hamPCR, see methods). Each point represents one leaf sample and is colored according to the leaf tissue: Green: ILZ, Purple: NIL, Orange, NILZ. A) Chao1 estimate of total alpha diversity, B) Shannon Index and C) Bacterial load in tissues from Ssc-infected plants (tan bars) compared to the control PDB treatment (grey bars). P-values are the result of a t-test comparing the treatments. D) and E) Constrained analysis of principle coordinates showing the effect of tissue type on the bacterial community structure, based on bray-curtis distances. PERMANOVA values reflect the result of 1000 permutations of the data.

### 3.3 SaxA affects bacterial diversity in the ILZ even in the absence of aliphatic glucosinolates

To test whether the observed patterns can be explained by detoxification of aITCs alone, we performed infections on the *A. thaliana* Col-0 *myb28/29* mutant, which is completely impaired in the biosynthesis of aGLSs. If aITCs were the main factor, we would expect to see strong bacterial enrichment in lesions of both Ssc WT and Ssc Δ*SaxA*. During infection of Col-0 *myb28/29* with Ssc WT there was no significant effect in the ILZ or in systemic tissue on either bacterial diversity or loads (Figure 4A-C). There was, however, a significant effect of the infection on beta diversity, with tissue type explaining about 15.8% of variation in the bacterial community during Ssc WT infection (Figure 4D and 4E, PERMAVONA, p = 0.007). In total 16 bacterial genera were differentially abundant between the ILZ and the NILZ (p < 0.001), all of which were higher abundance in the ILZ (Figure S5 and Supplementary Table 1). 5 of these 16 genera (*Pseudomonas, Simplicispira, Limnobacter, Rhizobacter* and *Marmoricola*) were also differentially abundant in the Col-0 WT/Ssc WT infection, but in the myb28/29 mutant the enrichment was less strong (log2FC < 4.5). Effects of PDB were again completely different from infection (Figure 4D and Figure S6). In contrast, lesions of Col-0 *myb28/29* infected with Ssc Δ*SaxA* showed a small decrease in alpha diversity but no significant effect on bacterial loads (Figure S7A-S7C) and no significant effect on beta diversity (Figure S7D and S7E, p = 0.43). Accordingly, no bacterial genera were significantly differentially enriched between the ILZ and NILZ below p < 0.05 (Figure S8 and Supplementary Table 1). Taken together, our data are consistent with a scenario in which fungal detoxification of aITCs in necrotic lesions by SaxA plays a positive role in shaping bacterial communities, since effects of Ssc WT infection were stronger in Col-0 WT than in Col-0 *myb28/29*. Some taxa appear to be consistently recruited to these lesions compared to surrounding tissue. On the other hand, SaxA was still relevant in the Col-0 *myb28/29* background, where aITCs cannot be produced, suggesting it has effects beyond aITCs.

**Figure 4:**
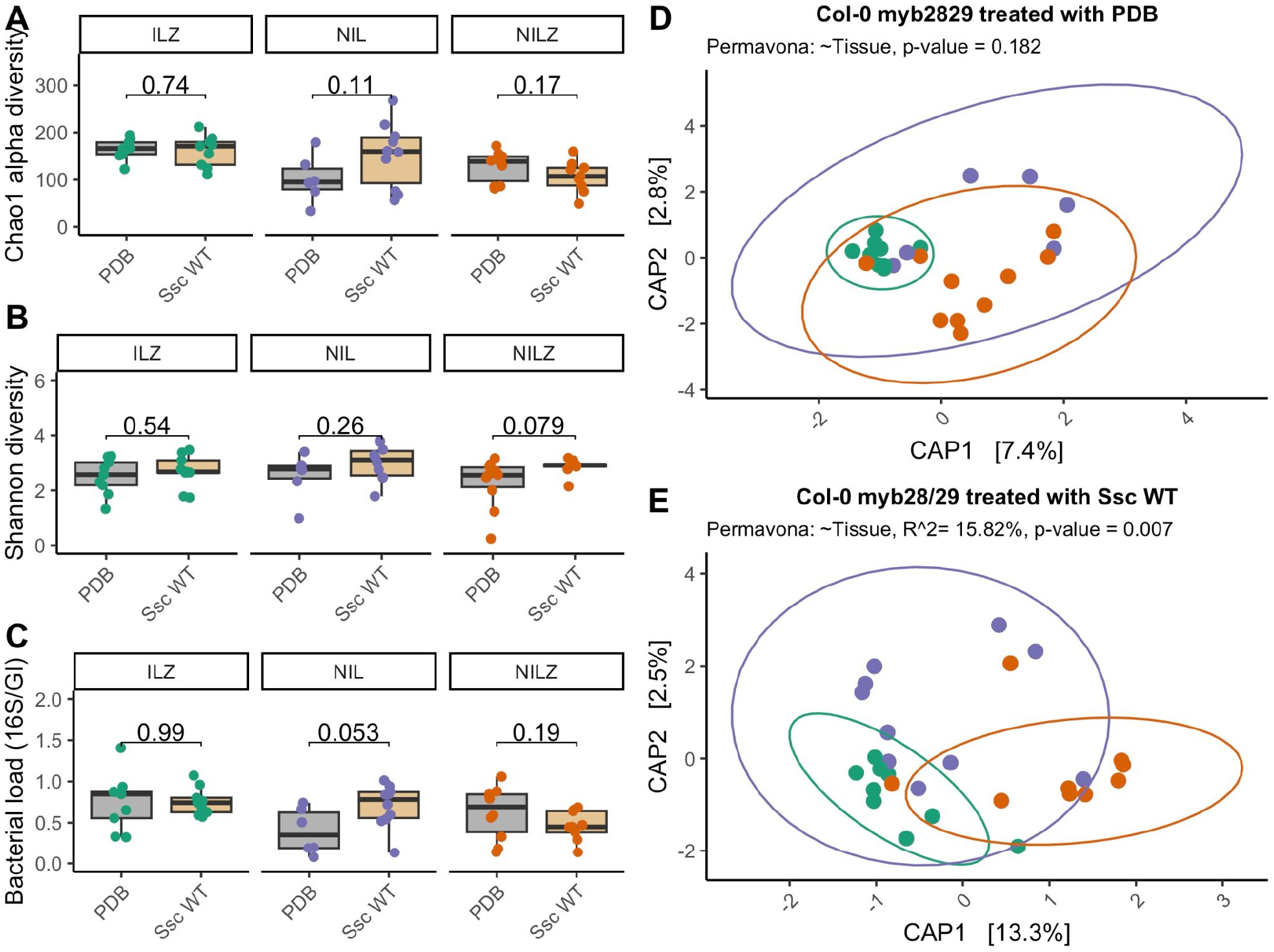
Bacterial community analysis of *A. thaliana* Col-0 *myb28/29* genotype infected with *S. sclerotiorum* WT. Measurements of leaf bacterial community composition of different leaf tissues (ILZ, NILZ and NIL - Defined in Figure 1) infected with Ssc *WT*. Alpha diversity results are based on 16S rRNA gene amplicon sequencing data, grouped at the genus level, while load and beta diversity results are the same data scaled to reflect absolute abundance (hamPCR, see methods). Each point represents one leaf sample and is colored according to the leaf tissue: Green: ILZ, Purple: NIL, Orange, NILZ. A) Chao1 estimate of total alpha diversity, B) Shannon Index and C) Bacterial load in tissues from Ssc-infected plants (tan bars) compared to the control PDB treatment (grey bars). P-values are the result of a t-test comparing the treatments. D) and E) Constrained analysis of principle coordinates showing the effect of tissue type on the bacterial community structure, based on bray-curtis distances. PERMANOVA values reflect the result of 1000 permutations of the data.

## 4. Discussion

Necrotophic fungal plant pathogens are highly destructive crop pests, causing massive crop losses under conducive conditions. Besides financial repercussions, the large-scale application of pesticides to control them also has far-reaching ecological consequences. Therefore, there is a strong incentive to develop alternative, more sustainable pathogen control strategies. *Sclerotinia sclerotiorum* (Ssc) has emerged as a model necrotrophic plant pathogen, in part due to its wide host range, including important crops. It has been suggested that bacterial biocontrol could be developed as an effective sustainable control strategy of Ssc (Han et al., 2023; Mülner et al., 2019). Most biocontrol discovery strategies, however, still rely on time-intensive large-scale screens that produce many false positives. Therefore, a better understanding of the mechanisms of Ssc-bacterial interactions is thought to be critical for more effective and rapid development (Wang et al., 2024).

We find that Ssc infection strongly interacts with leaf bacterial diversity in the model plant *Arabidopsis thaliana*. Effects of Ssc infection on the microbiome has not previously been thoroughly studied, although diseased lettuce stems were reported to have a relatively high amount of Gammaproteobacteria (Zhang et al., 2024). Changes to the host leaf microbiome caused by other eukaryotic pathogens has been well-documented. Some biotrophic oomycete and fungal leaf pathogens, for example, elicit large-scale changes on the host bacteriome of both infected (Agler et al., 2016; Jakuschkin et al., 2016) and systemic tissues (Goossens et al., 2023). Microbiomes of plants infected by necrotrophic pathogens are generally less well-studied. In *Zymoseptoria tritici*, the wheat leaf blotch pathogen, infection reshaped microbiomes in both infected and systemic tissue, mainly in resistant wheat cultivars (Seybold et al., 2020). Leaf infection with *Botrytis cinerea*, which is closely related to Ssc and has a similar lifestyle, induced microbiome changes in systemic rhizosphere tissues in tomato and strawberry (De Tender et al., 2016; Rizaludin et al., 2025), but local tissues were not investigated. While we did not investigate the rhizosphere microbiome, we did not detect effects in systemic leaf tissues, which may suggest that the most profound effects of SSc infection are local.

Our results indicate that aliphatic glucosinolates (aGLS) play an important role in determining the structure of the bacteriome of Ssc-induced necrotic lesions. This is consistent with previous work that suggests that remodelling and suppression of host immune metabolites by pathogens is an important mechanism shaping microbiomes. For example, infection of the obligate biotrophic oomycete *Albugo candida* alters the synthesis of indole glucosinolates in Arabidopsis, which allows increased colonization of non-adapted pathogens (Prince et al., 2017). Effects of the wheat leaf blotch pathogen *Zymoseptoria tritici* on local and systemic bacteriomes of resistant wheat cultivars were also correlated to strong changes to benzoxazinoids (BXs), metabolites that function similar to glucosinolates as phytoavengins (Kliebenstein and Kvitko, 2023; Seybold et al., 2020). In particular, upon infection, a susceptible cultivar showed no changes to glycosylated BX pools and limited effects on the bacterial microbiome, while BXs in a resistant cultivar shifted to the free, toxic form, together with an altered microbiome (Seybold et al., 2020). Our results similarly suggest that the toxic breakdown product of aGLS, aliphatic isothiocyanates (aITCs), were important in microbiome remodelling. However, in this case, it appears that ITC *detoxification* by SSc allows bacteria to grow in necrotic tissue, while in the *Z. tritici* example the presence of the toxic breakdown product most likely altered the microbiomes. The difference could be due to differences in the effects of the breakdown products or due to other host traits, or both. At any rate, a positive effect of detoxification on bacterial growth is consistent with previous work showing in-vitro that growth suppression of commensal bacteria by aITCs can be relieved by ITC detoxification by *Pseudomonas viridiflava* SaxA (Unger et al., 2025).

Interestingly, we observed that the fungal SaxA had a significant effect in both wild-type plants and in *myb28/29* mutant plants, which do not produce aGLS and therefore cannot have aITCs. So far, SaxA has been characterized strictly as an isothiocyanate hydrolase and other substrates are not known. Thus, a likely explanation is that SaxA also plays a role in detoxifying indole ITCs, derived from indole GLS. Indole GLS are, in contrast to constitutively produced aGLS, largely induced upon infection. In addition, they are still produced in *myb28/29* plants because they are derived from tryptophan in a pathway independent of aGLS (Sønderby et al., 2007). Indeed, during Ssc infection the indole GLS pool decreases in infected tissue, consistent with their hydrolysis to ITCs (Chen et al., 2021). Ssc SaxA attacks both aliphatic and aromatic ITCs, but indolic ITCs have not to our knowledge been tested (Chen et al., 2020). Thus, we hypothesize that the effect of SaxA on bacteria in necrotic lesions is shaped by both aliphatic and indole GLS and ITCs.

Ssc infection in *A. thaliana* Col-0 was also reported to cause accumulation of other anti-fungal compounds and camalexin, among other plant metabolites. Interestingly, however, we found only weak effects on bacteria in necrotic lesions in the Col-0 *myb2829* / Ssc Δ*SaxA* system compared to healthy tissue. The expected release of nutrients and the lack of plant immunity in necrotic tissue, as well as the lack of aliphatic glucosinolate-related metabolites might be expected to greatly increase bacterial colonization in these tissues. One explanation could be that Ssc produces antibiotics or antimicrobial peptides. For example, both oomycete and fungal plant pathogens have been shown to produce antimicrobial effector proteins that target specific bacteria to promote virulence (Gómez-Pérez et al., 2023; Snelders et al., 2021). Ssc secretes a range of effector proteins, but most of those characterized so far are only known to induce necrosis in hosts (Newman et al., 2023) and none have so far been described to affect plant microbiomes. If this were the case, however, we would expect to observe bacterial *suppression* in lesions in WT plants as well, where we mostly observed enrichment. While we cannot exclude antimicrobial production, our results are more consistent with the lack of microbiome effects in Ssc Δ*SaxA* lesions being due to accumulation of ITCs (aliphatic and indolic in WT plants and indolic in *myb28/29* plants). ITCs are toxic to a range of commensal leaf bacteria (Unger et al., 2024), and therefore may function as an efficient mechanism of clearing opportunistic bacteria from wounds, which could help prevent opportunistic infections.

In conclusion, the broad-range necrotrophic fungal pathogen Ssc strongly rearranges the host microbiome in necrotic lesions. This rearrangement is almost completely dependent on detoxification of ITCs, the breakdown products of glucosinolates, demonstrating that interactions of host and pathogen factors together shape the bacteriome. It is unclear what role this rearrangement plays on the outcome of Ssc infection. However, bacteria recruited to leaves during pathogen infection can in some cases contribute to reduced fungal virulence (Goossens et al., 2023). Thus, it is possible that Ssc bacteria may reduce Ssc virulence, for example via nutrient competition, or that they could increase virulence, for example by further inducing alterations to immune signals in neighboring tissues. Thus, future work should focus on elucidating these effects.

## Acknowledgements

This work was funded by the Deutsche Forschungsgemeinschaft (DFG, German Research Foundation) under under Germany’s Excellence Strategy - EXC 2051 - Projektnummer 390713860 (MTA) as well as under Projektnummer 458884166 (MTA | SS). It was also supported by the Max Planck Society (JC).

We further thank Dr. Daniel Vassao and Prof. Dr. Jonathan Gershenson for approving the use of the *S. sclerotiorum* strains.

## Data availability

All raw sequencing data will be made available in a public repository prior to peer-reviewed publication. Processed data and R code to generate the figures and supplementary table are being made publicly available at Figshare and are immediately available upon request from the authors.

## Notes

### Competing Interest Statement

The authors have declared no competing interest.

