## Supplementary Information for "Plant defense metabolite detoxification by a pathogenic fungus opens necrotic tissue to diverse bacteria in *Arabidopsis thaliana*"

Supplementary Figure S1-S8

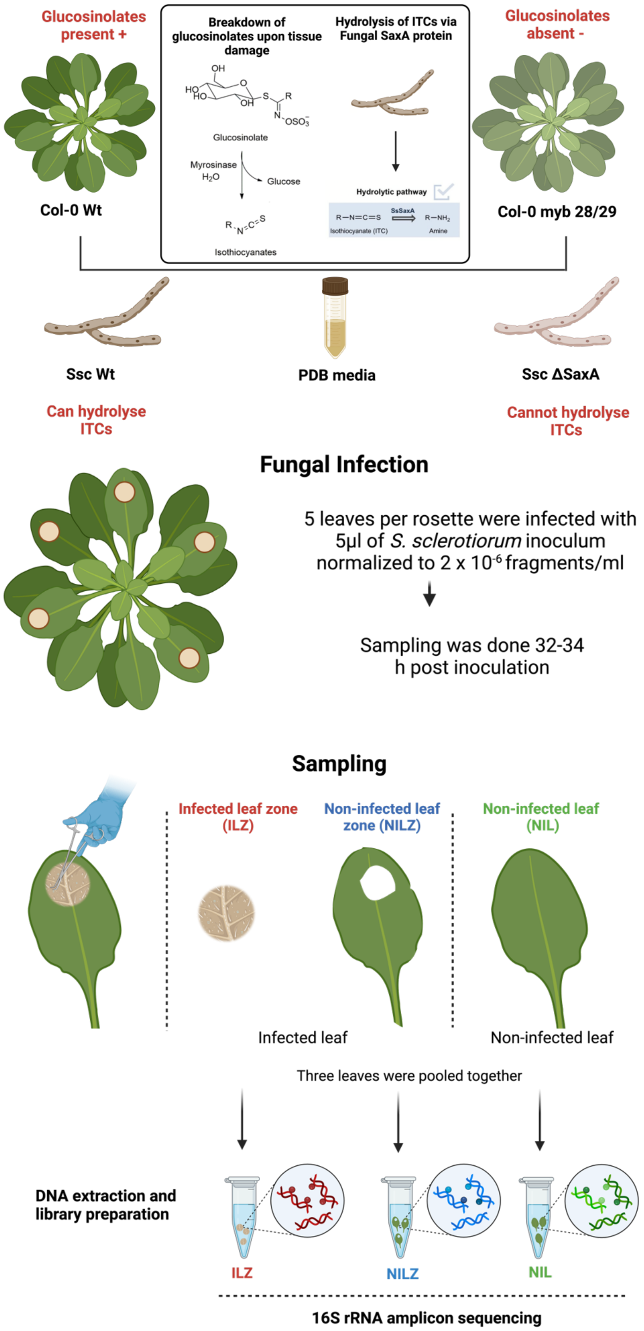


**Figure S1: Detailed diagram of the experimental approach for investigating bacterial communities asssociated with *S. sclerotiorum*-infected *A. thaliana* leaves.** A. thaliana Col-0 and myb28/29 mutant plants grown in a natural soil-supplemented substrate were drop-inoculated with Ssc WT, Ssc Δ*SaxA* or a PDB control. Leaves inoculated with Ssc developed necrotic lesions at the site of inoculation.

Inoculated leaves were dissected so that bacterial communities could be characterized in necrotic tissue (ILZ), non-necrotic tissue of the same leaf (NILZ) and in systemic tissue (NIL). PDB-inoculated control plants were dissected similarly. The leaves were then subjected to 16S rRNA gene amplicon sequencing for bacterial community analyses.

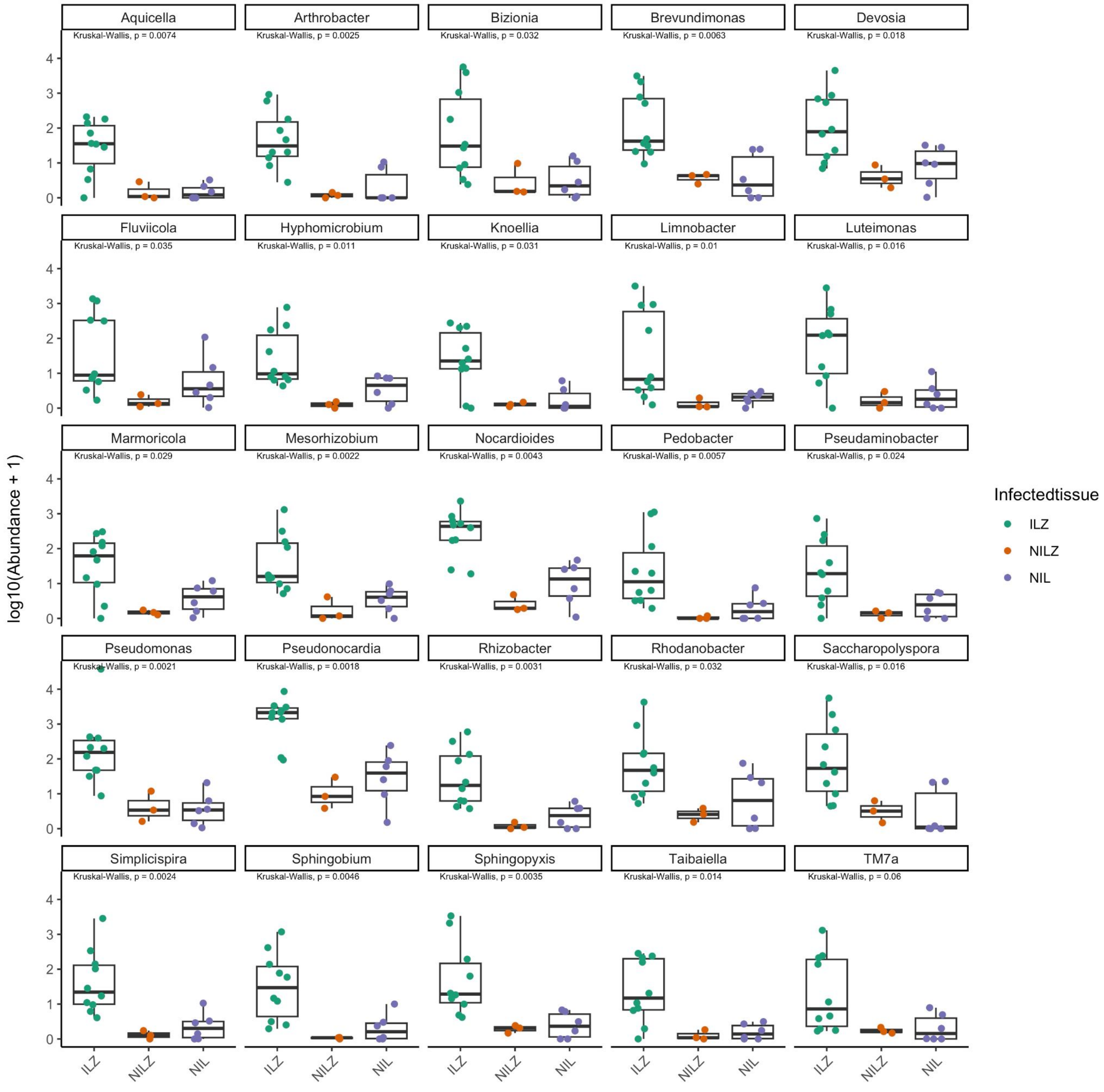


**Figure S2: Differentially abundant taxa between different leaf tissues of *S. sclerotiorum* WT-infected *A. thaliana* Col-0 plants.** DESeq2 analysis based on genus-level 16S rRNA amplicon sequencing data, scaled to reflect absolute abundance (hamPCR, see methods). A p-value cutoff of alpha = 0.001 was used to recover genera predicted to be differentially abundant taxa between the ILZ and NILZ. NIL is also plotted for comparison. Abundances of significant taxa were log10-transformed and plotted using an additional Kruskal-Wallis test for differences between the tissues.

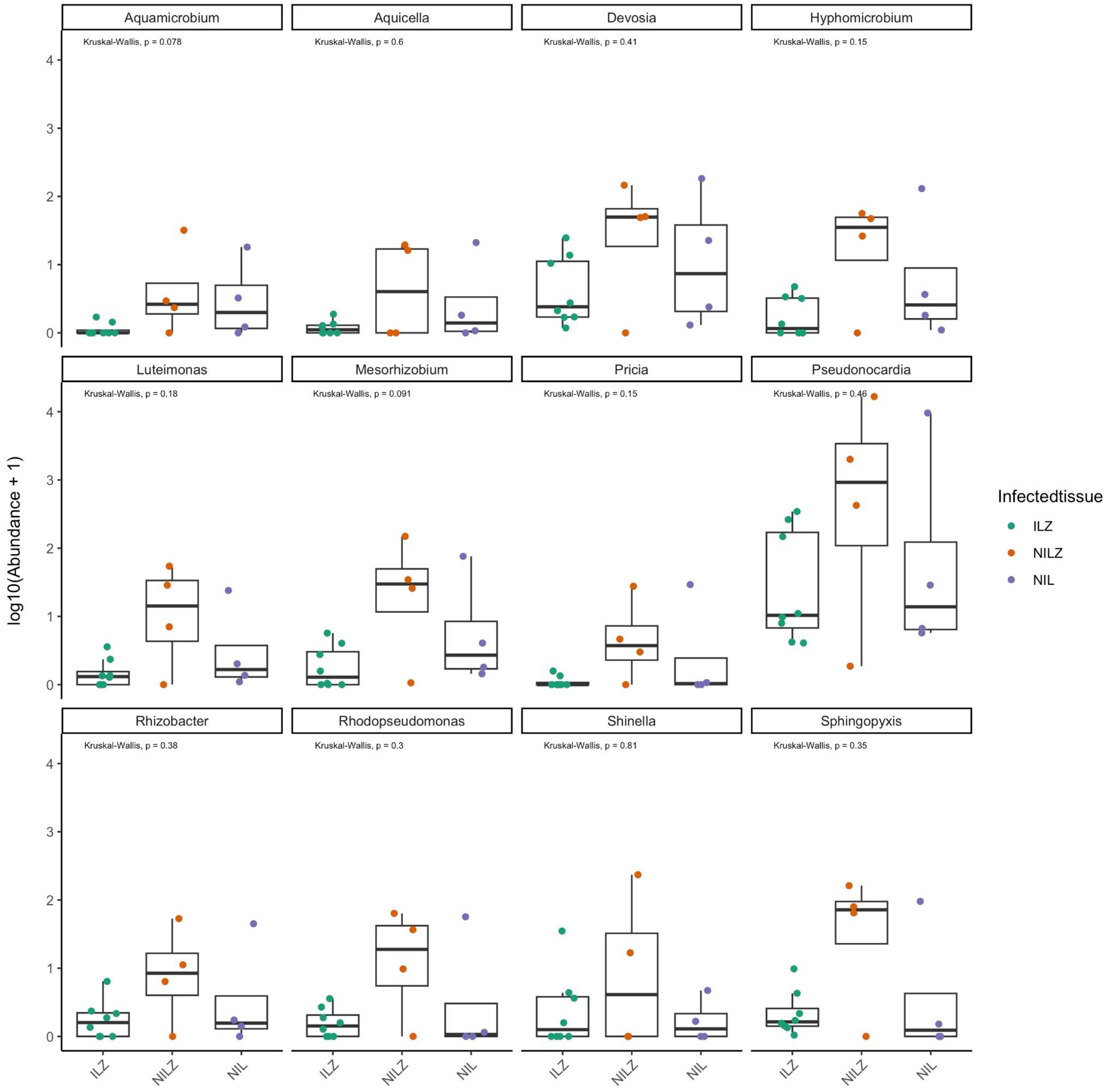


**Figure S3: Differentially abundant taxa between different leaf tissues of control PDB-treated *A. thaliana* Col-0 plants.** DESeq2 analysis based on genus-level 16S rRNA amplicon sequencing data, scaled to reflect absolute abundance (hamPCR, see methods). A p-value cutoff of alpha = 0.05 was used to recover genera predicted to be differentially abundant taxa between the ILZ and NILZ. NIL is also plotted for comparison. Abundances of significant taxa were log10-transformed and plotted using an additional Kruskal-Wallis test for differences between the tissues.

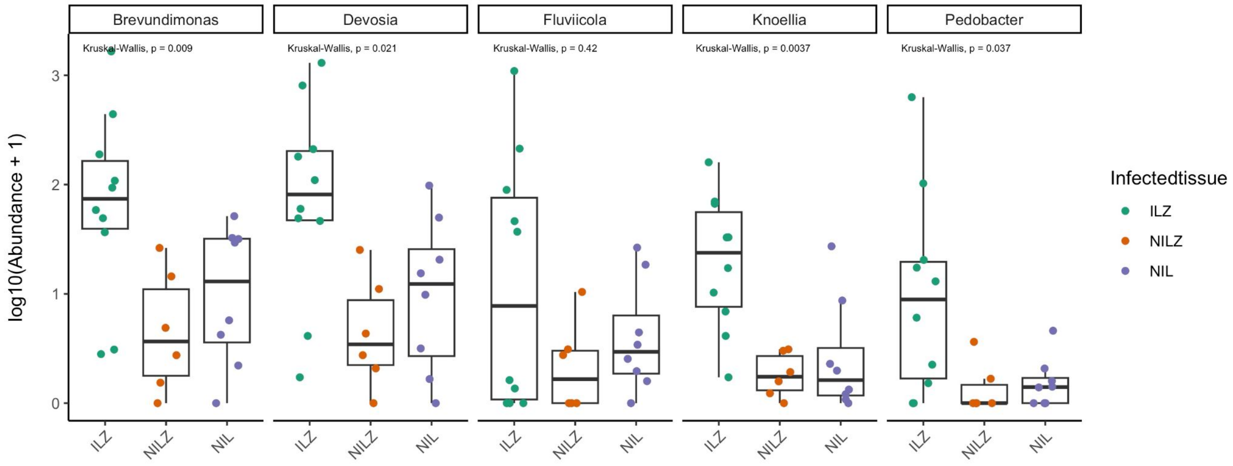


**Figure S4: Differentially abundant taxa between different leaf tissues of *S. sclerotiorum* Δ*SaxA*-infected *A. thaliana* Col-0 plants.** DESeq2 analysis based on genus-level 16S rRNA amplicon sequencing data, scaled to reflect absolute abundance (hamPCR, see methods). A p-value cutoff of alpha = 0.001 was used to recover genera predicted to be differentially abundant taxa between the ILZ and NILZ. NIL is also plotted for comparison. Abundances of significant taxa were log10-transformed and plotted using an additional Kruskal-Wallis test for differences between the tissues.

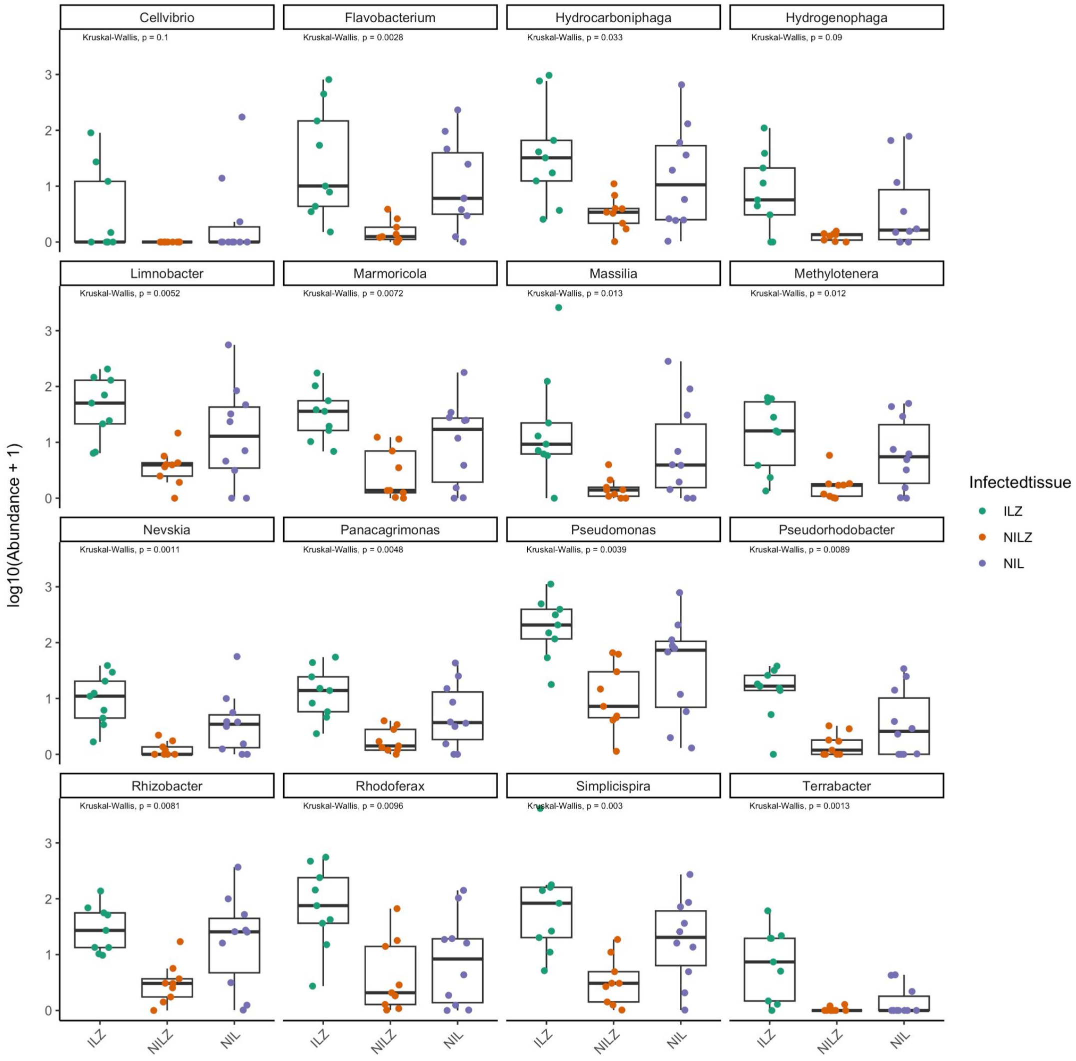


**Figure S5: Differentially abundant taxa between different leaf tissues of *S. sclerotiorum* WT-infected *A. thaliana* Col-0 *myb28/29* plants.** DESeq2 analysis based on genus-level 16S rRNA amplicon sequencing data, scaled to reflect absolute abundance (hamPCR, see methods). A p-value cutoff of alpha = 0.001 was used to recover genera predicted to be differentially abundant taxa between the ILZ and NILZ. NIL is also plotted for comparison. Abundances of significant taxa were log10-transformed and plotted using an additional Kruskal-Wallis test for differences between the tissues.

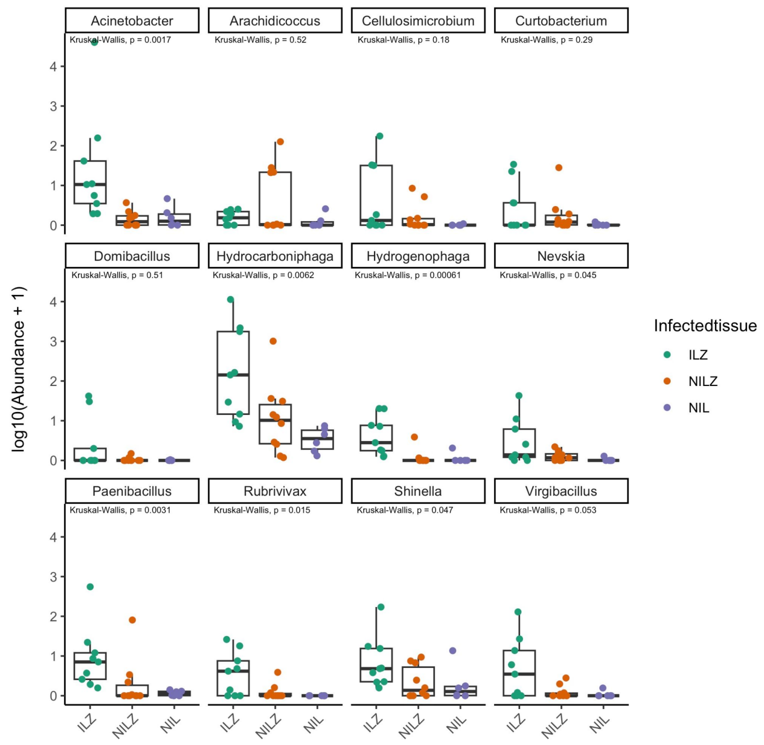


**Figure S6: Differentially abundant taxa between different leaf tissues of control PDB-treated *A. thaliana* Col-0 *myb28/29* plants.** DESeq2 analysis based on genus-level 16S rRNA amplicon sequencing data, scaled to reflect absolute abundance (hamPCR, see methods). A p-value cutoff of alpha = 0.05 was used to recover genera predicted to be differentially abundant taxa between the ILZ and NILZ. NIL is also plotted for comparison. Abundances of significant taxa were log10-transformed and plotted using an additional Kruskal-Wallis test for differences between the tissues.

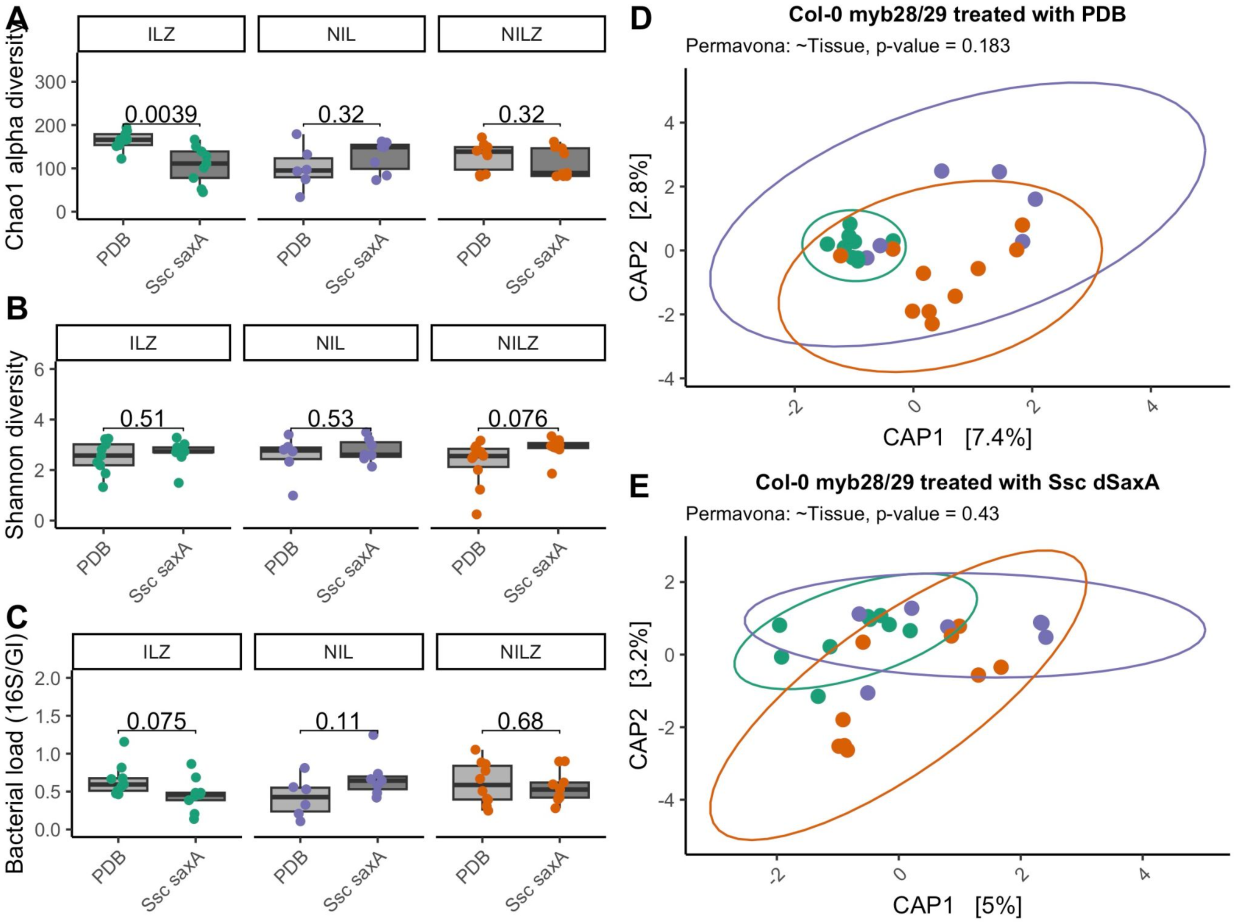


**Figure S7: Bacterial community analysis of *A. thaliana* Col-0 *myb28/29* genotype infected with *S. sclerotiorum* Δ*SaxA***. Measurements of leaf bacterial community composition of different leaf tissues (ILZ, NILZ and NIL - Defined in Figure 1) infected with Ssc Δ*SaxA.* Alpha diversity results are based on 16S rRNA gene amplicon sequencing data, grouped at the genus level, while load and beta diversity results are the same data scaled to reflect absolute abundance (hamPCR, see methods). Each point represents one leaf sample and is colored according to the leaf tissue: Green: ILZ, Purple: NIL, Orange, NILZ. A) Chao1 estimate of total alpha diversity, B) Shannon Index and C) Bacterial load in tissues from Ssc-infected plants (tan bars) compared to the control PDB treatment (grey bars). P-values are the result of a t-test comparing the treatments. D) and E) Constrained analysis of principle coordinates showing the effect of tissue type on the bacterial community structure, based on bray-curtis distances. PERMANOVA values reflect the result of 1000 permutations of the data.

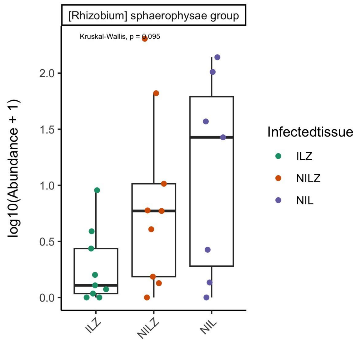


**Figure S8: Differentially abundant taxa between different leaf tissues of *S. sclerotiorum* Δ*SaxA*-infected *A. thaliana* Col-0 *myb28/29* plants.** DESeq2 analysis based on genus-level 16S rRNA amplicon sequencing data, scaled to reflect absolute abundance (hamPCR, see methods). A p-value cutoff of alpha = 0.1 was used to recover genera predicted to be differentially abundant taxa between the ILZ and NILZ. NIL is also plotted for comparison. Abundances of significant taxa were log10-transformed and plotted using an additional Kruskal-Wallis test for differences between the tissues.
